# Focused small field of view magnetic particle imaging for the detection and quantification of tumour associated macrophages

**DOI:** 10.1101/2023.06.22.545970

**Authors:** Nitara Fernando, Julia J. Gevaert, Justin Konkle, Patrick Goodwill, Paula J. Foster

**Author notes:** Corresponding Author: Nitara Fernando, Department of Medical Biophysics, Robarts Research Institute, University of Western Ontario, 1151 Richmond Street North, London, Ontario N6A 5B7.

## Abstract

Tumour Associated Macrophages (TAMs) play a crucial role in breast cancer progression and have the potential to be used as a biomarker for patient prognosis. Magnetic particle imaging (MPI) is an emerging modality which can detect cells labelled with superparamagnetic iron oxide (SPIO) nanoparticles and can be used for non-invasive TAM assessment. However, MPI TAM detection is limited by its effective dynamic range. This limitation occurs when SPIO nanoparticles injected intravenously accumulate in the liver resulting in a large MPI signal which shadows regions of interest with lower signals (i.e the tumour) preventing their isolation and quantification. In this study we test an advanced reconstruction algorithm which allows us to prescribe a small focused field of view (FOV) on lower signals of interest. We then demonstrate the success of this method with an in vivo tumour model and show enhanced image quality and successful quantification of TAMs in mouse mammary tumours with different metastatic potentials (4T1 and E0771). Utilizing in vivo MPI, we did not see significant differences in the MPI signal for 4T1 tumours compared to E0771. These findings highlight the potential of MPI for in vivo TAM quantification offering a promising avenue for broader applications in cancer research and potentially overcoming constraints of MPI in other in vivo imaging contexts.

## Introduction

Tumour associated macrophages (TAMs) are a highly prevalent component of the tumour microenvironment (TME), and they can constitute up to 50% of the breast cancer TME^1^. In breast cancer, higher TAM infiltration has been associated with poorer patient prognosis^1-3^. TAM density can be assessed using immunohistochemistry (IHC); however, this requires invasive biopsies and is not representative of the whole tumour^4^. Thus, there is a need for non-invasive and quantitative imaging that allows for the in vivo assessment of TAMs, which could serve as a biomarker for tumour aggressiveness.

Superparamagnetic iron oxide (SPIO) particles can be used to label macrophages in situ via an intravenous (IV) injection and this approach has been used for imaging TAMs with MRI^5-7^. However, quantification of TAMs from iron-induced signal loss in MRI is challenging. Magnetic Particle Imaging (MPI) is emerging as an in vivo cellular imaging method and can be an alternative to MRI for cell tracking^8^. MPI directly detects the nonlinear magnetization of superparamagnetic iron oxide (SPIO) particles as a signal hot-spot, as opposed to MRI which indirectly detects SPIO as regions of signal loss from its effect on proton relaxation. The MPI signal is quantifiable as it is linearly proportional to iron mass; with knowledge of the cellular iron loading, the number of cells can be estimated.

MPI has previously been evaluated for the detection and quantification of TAMs, however, the high uptake of SPIO in liver macrophages after IV injection prevented isolation of the lower TAM signal in vivo. This is due to known dynamic range limitations in MPI^9,10^; when iron samples with large differences in concentrations are present in the same field of view (FOV) there is signal oversaturation from the higher signal due to the requirement for regularization for stable reconstruction. This represents a major roadblock for in vivo MPI in applications where two or more sources of signal exist.

In this study we addressed this challenge by using a small FOV focused on the tumour, together with an advanced reconstruction algorithm. In 2015, Konkle et al. introduced an advanced reconstruction method which employs a priori information, non-negativity, and image smoothness to enhance image quality^11^. First, we evaluated this using samples of iron, where a sample with a high iron concentration was positioned 2 cm away from a sample with a much lower iron concentration and showed that this approach removed image artifacts seen with the native MPI reconstruction. Then, we employed this strategy for quantitative imaging of tumours in vivo after IV SPIO and compared the MPI signal for mouse mammary tumours induced using two breast cancer cell lines with different metastatic potentials.

## Methods and materials

### MPI System and Image Reconstruction

All experiments were performed using the Momentum™ pre-clinical MPI scanner (Magnetic Insight Inc.). The Momentum scanner is an oil-cooled, field free line (FFL) scanner that uses an alternating magnetic field to excite SPIO particles and reconstructs images using x-space signal processing^12^. The native image reconstruction is the default reconstruction method available in the MOMENTUM regular user interface (RUI) and is formed by the X-space stitching method^11,13^. In this method the panels of data are stitched together, and the edges of the image are pinned to zero, which allows for the recovery of the filtered DC component. To acquire the images using the advanced reconstruction, we enabled the ‘inverse_xz_image_combiner’ option in the MOMENTUM Advanced User Interface (AUI). This option reconstructs native images using x-space methods as described above and applies an inverse problem postprocessing step. The postprocessing combines native images that were reconstructed separately depending on the direction of transmit (positive and negative), sharpens using the equalized PSF^14^, and corrects the edge-pinning applied in the native image reconstruction step. The postprocessing is formulated as a matrix-free non-negative least squares inverse problem and is solved using FISTA as in previous formulations^11^.

### Magnetic Particles

Synomag-D (Micromod GmbH, Germany) was the SPIO used for all in vitro dynamic range experiments. Synomag-D is a multicore particle with a nanoflower substructure^15^. The agglomerated iron core size is ∼30 nm, coated with dextran to improve stability and provide compatibility. The hydrodynamic diameter was ∼50 nm. For the in vivo tumour model experiments, a polyethylene glycol (PEG) coated version of Synomag-D was used (Synomag-D PEG 25.000-OMe). The hydrodynamic diameter is ∼70nm.

### In Vitro: Idealized Dynamic Range – Single Sample – Full FOV

The idealized dynamic range of the MPI system was tested by measuring samples of different concentrations individually. A 1:1 dilution series was prepared using known amounts of the SPIO, Synomag-D. Twelve samples were prepared ranging from 50.0 µg Fe to 24.4 ng Fe in 5 µL phosphate buffer solution (PBS) (Table 1).

**Table 1.**
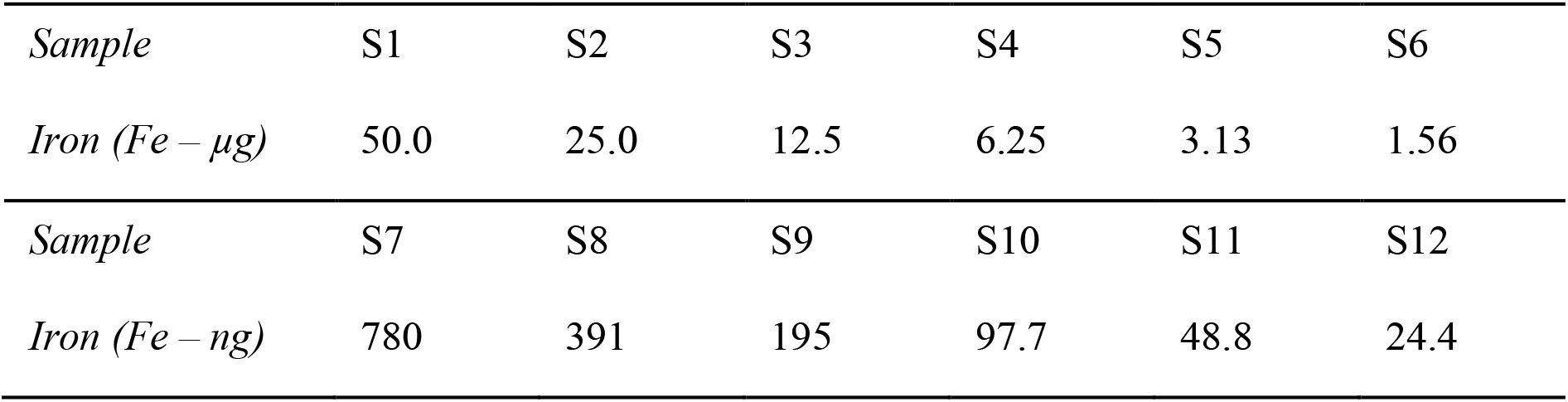
Sample preparation for dynamic range experiments.

Samples were individually imaged in the center of a 6 × 6 × 12 cm (X, Y, Z) field of view (FOV). Projection images were acquired in 2D with a 3.0 T/m selection field gradient and drive field strengths of 20 mT and 23 mT in the X and Z axes, respectively, with an acquisition time of ∼2 minutes. These 12 cm FOV images were reconstructed using a native image reconstruction enabled in the RUI of the MPI system.

### In Vitro: Effective Dynamic Range – Two Samples - Full FOV

The effective dynamic range of the system was tested by measuring samples with different concentrations together within the same 12 cm FOV. In this experiment, S1 (50.0 µg Fe) was placed 2 cm apart from samples with decreasing iron concentrations (S2, S3, S4, etc.) (Figure 2A-H).

### In Vitro: Two Samples – Small FOV with advanced image reconstruction

In this experiment, S7 (780 ng Fe) sample was placed 2 cm apart from S1 (50 µg Fe) and MPI was repeated with triplicate samples. A 6 × 2 cm (X, Z) FOV was centered on S7 and images were acquired for each sample (n = 3) using the standard native reconstruction algorithm in the RUI and the advanced reconstruction algorithm enabled in the AUI.

### Cell Culture

E0771 and 4T1 murine breast cancer cell lines were utilized. Cells were cultured at 37 °C and 5% CO2. E0771 cells were maintained in RPMI (Gibco, Thermo Fisher Scientific, MA, USA), while 4T1 cells were cultured in DMEM (Gibco, Thermo Fisher Scientific, MA, USA). The culture media for both cell lines were supplemented with 10% fetal bovine serum and 1% antimycotic/antibiotic, and cells were passaged every 2–3 days. The 4T1 cell line is highly tumorigenic and invasive and can spontaneously metastasize from the primary tumour in the mammary fat pad to multiple distant sites including lymph nodes, blood, liver, lung, brain, and bone^16,17^. E0771 is poorly metastatic compared to 4T1^18^.

### In Vivo Tumour Imaging

BALB/c and C57BL/6 mice were purchased from Charles River Laboratories, Inc. (Senneville, CAN). All animal studies were performed in accordance with institutional and national guidelines. Mice were anesthetized with isoflurane administered at 2% in oxygen. Subsequently, 100, 000 4T1 and E0771 cells (>90% viability measured using the trypan blue exclusion assay), suspended in a 50 uL PBS solution, were administered subcutaneously to the fourth mammary fat pad of C57BL/6 mice (E0771, n = 11) and BALB/c mice (4T1, n = 11). Animals were monitored every other day and tumour volumes were measured using calipers.

Twenty days after the cell injection, 30mg/kg of Synomag-D-PEG was administered IV to mice in a 100 µL volume via tail vein to 8 mice in each group, the other 3 mice in each group were used as controls and did not receive SPIO. MPI imaging was performed 24 hours later. 2D and 3D tomographic images were acquired with a 5.7 T/m selection field gradient. Mice were first imaged with a 6 × 6 × 12 cm (X, Y, Z) FOV with the native reconstruction and multichannel images were acquired (drive field strengths: 23 mT (z) and 20 mT(x)). Next, mice were imaged with a 6 × 6 × 3 cm (X, Y, Z) FOV focused on the tumour region using the advanced reconstruction, and single channel images were acquired (drive field strength: 23mT (z)).

### Ex vivo tumour imaging

24 hours following the last MPI exam, mice were sacrificed. Tumours from mice that received SPIO and mice that did not receive SPIO (controls) were removed and placed in 4% formalin for 24 hours for fixation. Then, tumours were cryoprotected by passing through a sucrose gradient, with concentrations of 15% and 30%. Ex vivo MPI and MRI were performed on the cryoprotected tumours.

3D MPI images were acquired using the same parameters used for in vivo imaging (advanced reconstruction, 3 cm FOV, single channel).

### MPI Analysis and Quantification

All MPI images were imported into Horos™, an open-source clinically relevant image analysis software (version 3.3.6, Annapolis, MD USA). Images were viewed using a custom MPI colour look-up table (CLUT).

For in vitro analysis of MPI dynamic range, an image was acquired of the empty sample holder for each image FOV, to account for the background noise. The standard deviation of the background noise was measured from a region of interest (ROI) encompassing the entire background image and a threshold value of 5 times the standard deviation of the background noise was applied for subsequent image analysis to ensure only signal above this limit was measured. We have studied various image analysis methods for MPI and found this method to produce accurate quantification with low user variability^19^. Using this threshold, a semi-automatic segmentation tool was used to measure the mean MPI signal within a specific ROI.

For analysis of in vivo tumour signal, a threshold value of 0.5 times the maximum tumour signal was applied as a lower bound for signal detection^19^. This method was chosen as it allowed us to ensure no signal due to iron in the tail from IV injections was detected when quantifying tumour signal.

Total MPI signal for an ROI was calculated by multiplying the ROI area (2D) or volume (3D) by the mean signal. Calibration lines were made to show the relationship between iron mass and total MPI signal. Calibration lines were used to quantify iron mass from images. A separate calibration must be made for every different SPIO and parameter set. Iron mass was calculated by dividing the total MPI signal by the slope of the respective calibration line for those image parameters. All MPI images were segmented and analyzed in the same way to ensure consistency.

### Statistical Methods

Statistical analyses were performed using GraphPad Prism Software. A simple linear regression was used to determine the relationship between iron mass and total MPI signal for the 2 cm, 3 cm and 12 cm FOV calibration lines. These were used for subsequent iron quantification from measured total MPI signal in images. Deviations greater than 10% from known iron values were not considered accurate. For the effective dynamic range experiment (two samples, small FOV) an average of the iron mass estimated from triplicate measurements of S7 was used to compare to the known iron mass using a one sample t and Wilcoxon test. For the in vivo tumour signal analysis, a Mann-Whitney test was used to compare signal between the two cancer cell types; p>0.05 was considered a significant finding.

### Microscopy

Following cryoprotection, the tumour samples were embedded in an OCT compound, sectioned (10 μm thickness), and then stained with hematoxylin and eosin (H&E), Perls’ Prussian Blue (PPB) for iron, and a pan macrophage stain (CD 68). Microscopy was performed using the Echo 4 Revolve Microscope (CA, USA).

## Results

### In Vitro: Idealized Dynamic Range – Single Sample - Full FOV

The idealized dynamic range describes the range of detectable signal from varying iron concentrations when there is only one source of signal within the imaging FOV. Individually imaged samples were detected and quantified from 50 µg Fe (S1) to as low as 48.8 ng Fe (S11) with the chosen parameters. At the next lowest sample (S12, 24.4 ng Fe), the MPI signal was not higher than the threshold limit (5 times the standard deviation of the background signal) and was not considered quantifiable (Figure 1a). In the S12 image, “ghost” signal is observed in the background which is in the same range as the low iron sample. From the S1-S11 images, a calibration line relating MPI signal and iron mass was made and analyzed using a simple linear regression model. There was a strong correlation (R^2^ > 0.9) between MPI signal and iron mass. The slope of the calibration line was used for quantifying samples in subsequent effective dynamic range experiments (Figure 1b).

**Figure 1:**
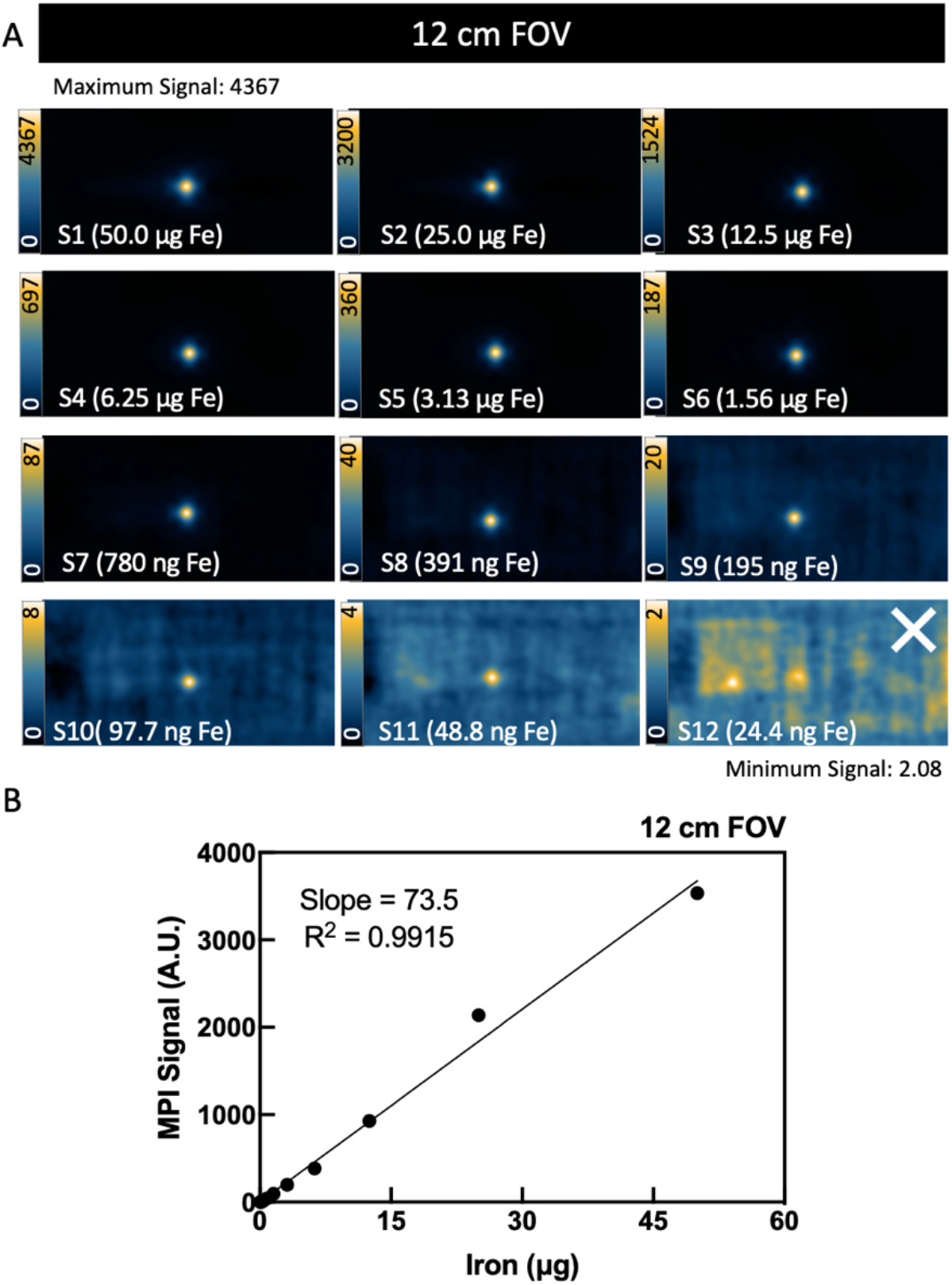
MPI idealized dynamic range for single samples and a full 12 cm FOV. (A) Individually imaged samples were detected and quantified until sample 11 (S11, 48.8 ng Fe) as the detection threshold. (B) Samples 1-11 were quantified and related to MPI signal, showing a strong linear correlation (R^2^ > 0.9), creating a calibration line used for downstream quantification. The signal from S12 was not detectable and was not quantified, indicated by the white X.

The lower bound of the idealized dynamic range was determined by considering the detection of S1-S11 but not S12, representing a dilution factor of 2^10^ = 1024. The upper bound was not tested beyond S1 (50 µg Fe), since iron concentrations above this amount approached system saturation levels based on MPI relaxometry testing. At very high iron concentrations, the interactions between iron particles tend towards non-linear relations between the particle concentration and the generated signal^20^.

### In Vitro: Effective Dynamic Range – Two Samples - Full FOV

The effective dynamic range describes the range of detectable signal when there are two or more signals of varying concentrations within the imaging FOV. In these experiments, a 2-sample model was used where a sample containing a high iron concentration (S1, 50 µg Fe) was placed 2 cm apart from a sample with a lower iron concentration (S2 – S8). Quantification of S1 was consistently within 10% of manufacturer reported values and considered accurate for effective dynamic range experiments. The two samples were imaged with a 12 cm FOV which encompassed both samples. At a 2 cm separation distance, the lower signals from S2-6 were resolved from the higher signal from S1, representing a detection limit of 2^6^ = 64. This limit describes the point at which the lower signal could be resolved from the higher signal using the described threshold method and quantified, though it does not account for accuracy of quantification. Quantification of S6 was considered inaccurate based on our definition of a deviation greater than 10% from known iron values. Therefore, the quantification limit was 2^5^ = 32 (Figure 2). At this limit, with window levels set to the minimum and maximum value of the lower signal, image artifacts begin to appear and become more exaggerated as the iron concentrations decrease. The oversaturated higher signal extends towards the edge of the field of view, resulting in large inverted negative signal artifacts central to the image, affecting low signal isolation and quantification.

**Figure 2:**
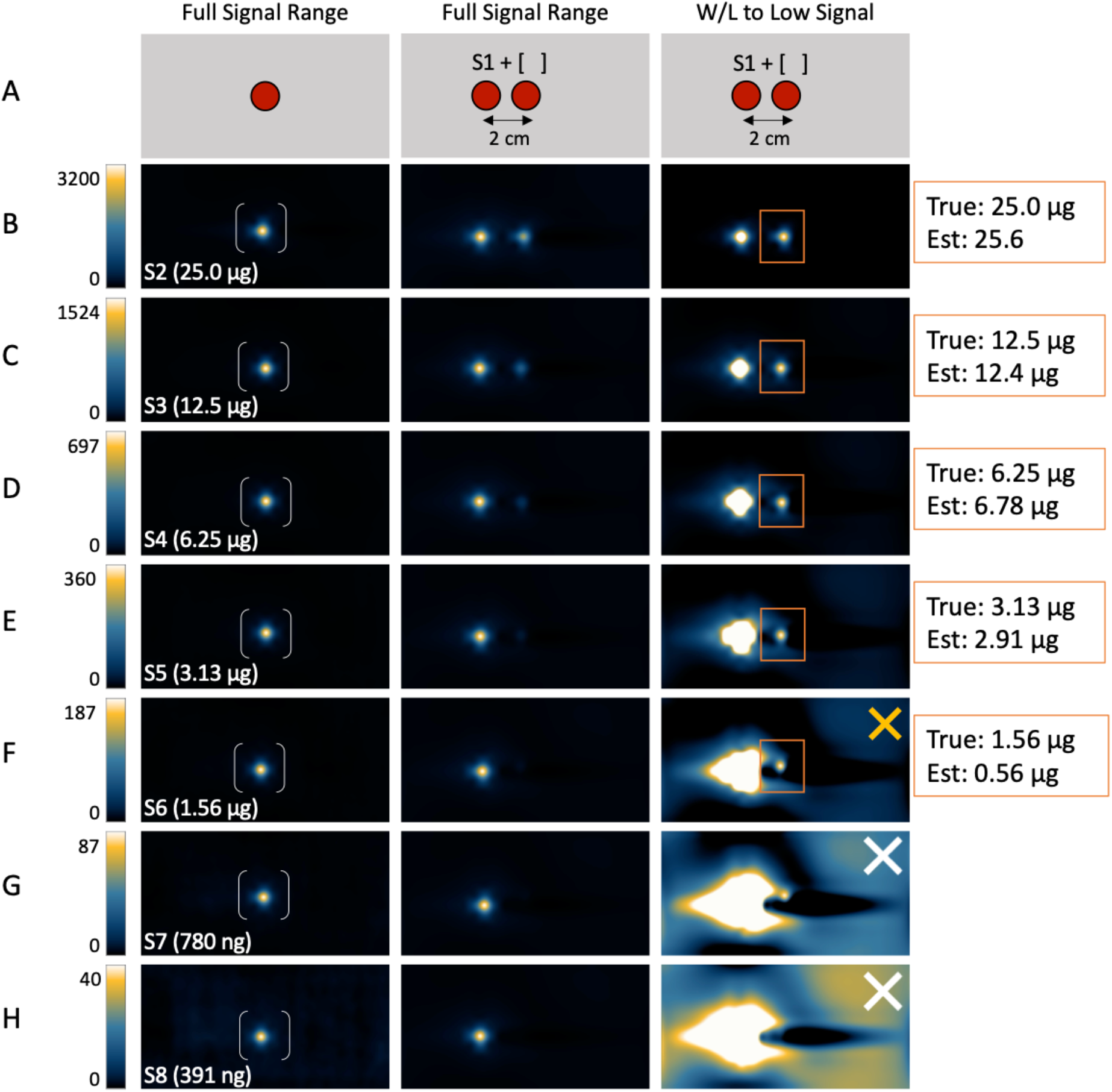
MPI effective dynamic range using two samples of varying iron mass at 2 cm separation imaged with a full 12 cm FOV. Here, the dynamic range of the system is limited to sample 6 (S6, 1.56 µg Fe), a much higher threshold than the idealized range from single sample imaging. Quantification begins to deviate more than 10% with S6. At the next lowest iron mass, S7, the signal cannot be distinguishably quantified from the much higher signal coming from S1 (50.0 µg Fe). In (f), the yellow X indicates signal that was detected but not accurately quantified. The white X in (g) and (h) indicates signal that was not detectable or quantifiable. True means iron mass from manufacturer reported values. Est. means estimated iron mass calculated from calibration lines.

### In Vitro: Two Samples – Small FOV with advanced image reconstruction

The purpose of this experiment was to investigate using a small, focused FOV to isolate and quantify proximal high and low signals. The same set-up as Figure 2 was used with a 2 cm separation distance between samples. S1 (50 µg Fe) was placed 2 cm apart from S7 (780 ng Fe) and imaged within a 12 cm FOV and the native reconstruction (Figure 3a). With the image shown in its full dynamic range (the minimum and maximum signal within the entire FOV), S1 has visible signal but S7 does not (Figure 3b – the same as Figure 2g, second image panel). Window leveling to the minimum and maximum signal of S7 resulted in oversaturation of the higher S1 signal, creating an inverted negative artifact and preventing quantification of the lower S7 signal (Figure 3c – the same as Figure 2g, third image panel). Using the same sample placement, a 2 cm FOV was centered on S7 (Figure 3d). Images were acquired using native reconstruction (Figure 3e) and advanced reconstruction (Figure 3f). Native reconstruction did not isolate S7 signal or resolve the image artifact which appeared as a negative signal. Native reconstruction automatically assumes there is no signal at the FOV edges and always sets these values to zero. This assumption causes an inverted negative signal artifact to appear on images when there is signal on the FOV edges that is not taken into reconstruction and is automatically set to zero. With this artifact present, quantification was not possible. The advanced reconstruction fully isolated the low signal within a 2 cm FOV with no image artifacts. Triplicates of S7 (780 ng Fe) were imaged this way with an average iron content of 797 ng Fe estimated from the calibration line. Estimated iron values for the triplicate samples were within 10% of the true value and there was no statistically significant difference between the estimated and true iron values (p > 0.05) (Figure 3g).

**Figure 3:**
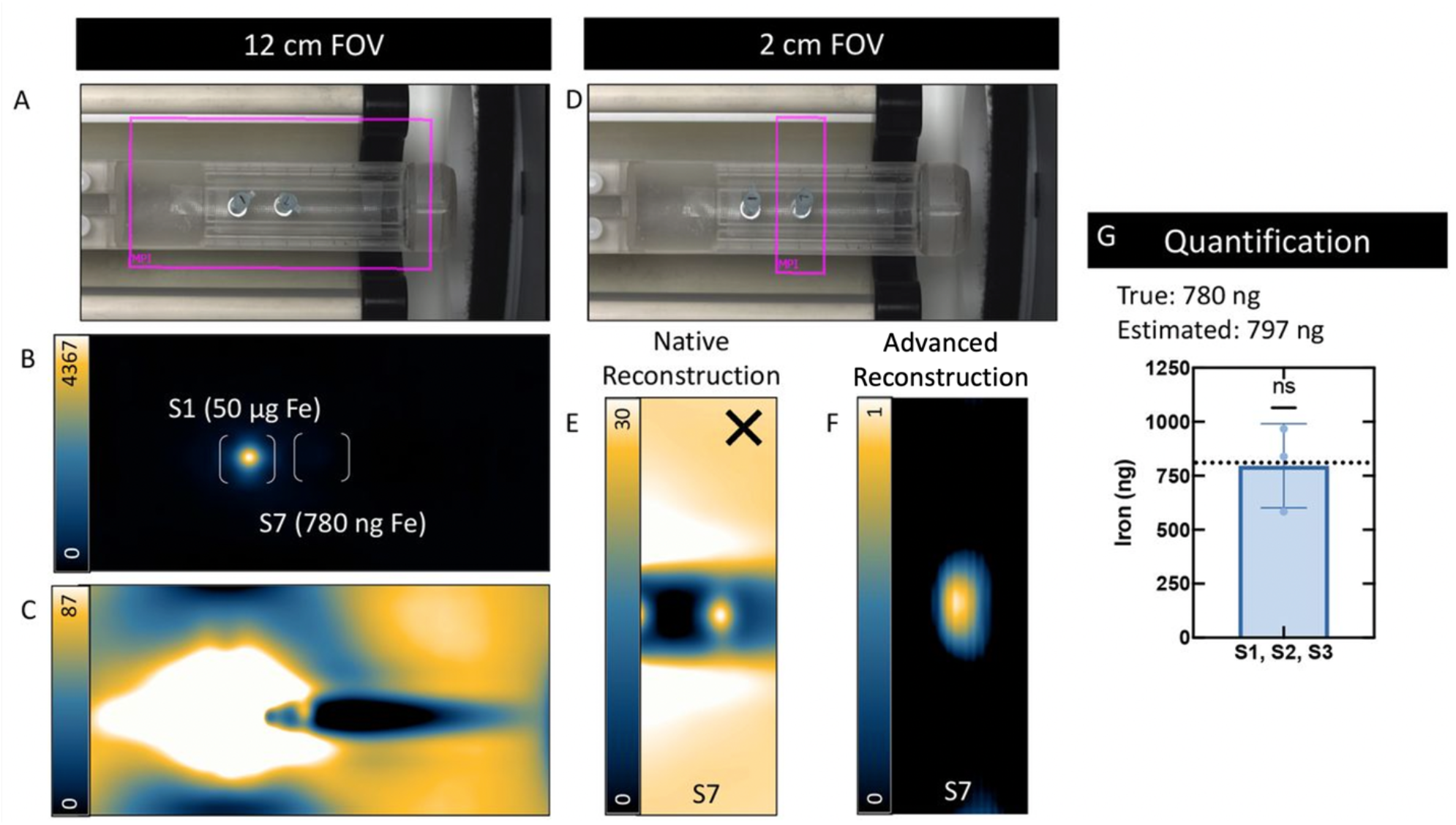
Small field of view (FOV) imaging with the advanced reconstruction as a solution to quantifying two proximal samples with varying iron mass. (a) Experimental set-up showing the full 12 cm FOV around a source of low signal (S7, 780 ng Fe) 2 cm apart from a higher signal (S1, 50 µg Fe). (b) Viewing the image in its full dynamic range only shows signal coming from the higher iron sample (S1). (c) Window leveling to the minimum and maximum signal of the lower signal (S7) oversaturates the high signal and prevents quantification of S7. (d) A 2 cm FOV was centered on the low sample (S7) and acquired with (e) native reconstruction, which did not quantify signal (indicated by the X) and (f) the advanced reconstruction, which did quantify signal. (g) The advanced reconstruction successfully isolated the low signal and triplicate samples were imaged with the average iron content estimated to be 797 ng Fe, in agreement with the known iron content (780 ng Fe) (p > 0.05).

### In Vivo Tumour Imaging

For imaging tumours with the full FOV (Fig. 4A) we observed signal oversaturation from the liver region of the mouse and were unable to isolate and quantify signal from the tumour. This is due to the previously described limitation in MPI dynamic range. This was solved using a FOV focused on the tumour region and the advanced reconstruction (Fig. 4B) and allowed MPI signal to be quantified for all tumours.

**Figure 4.**
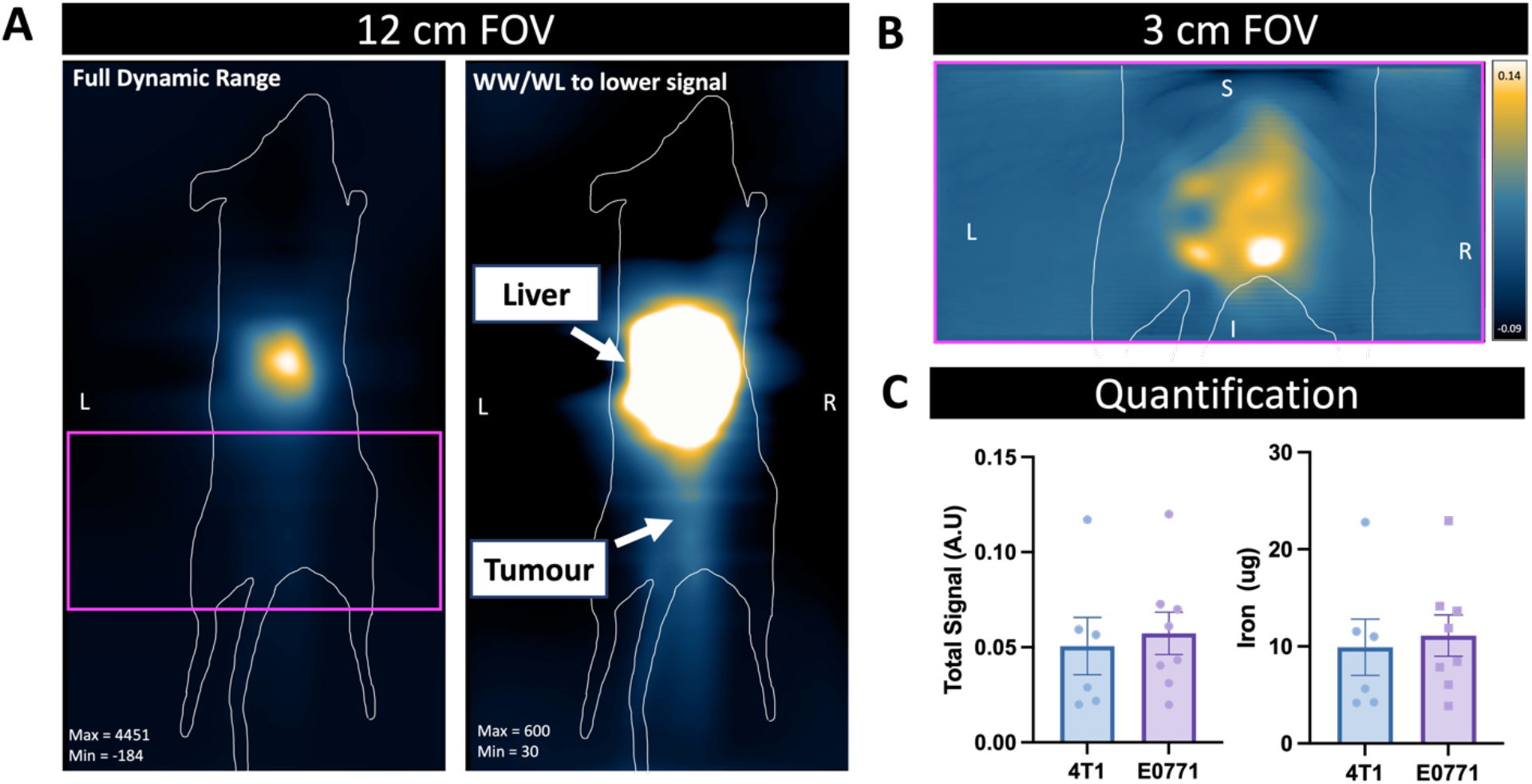
In vivo imaging of a representative E0771 tumour using MPI following tail vein injection of Synomag-D PEG. (A) The standard 12 cm FOV is used and the 2D MPI images are shown. Viewing the image in its full dynamic range, only the high liver signal is visible. Adjusting the window width and level to the maximum and minimum of the lower signal oversaturates the higher signal and interferes with the detection of the tumour signal. (B) A 3 cm FOV was applied over the tumour region, shown by the pink box, allowing for isolation of tumour signal. The signal obtained with small FOV imaging can be quantified (C) as total signal or iron content using a previously established calibration line.

MPI signal and iron content were quantified for 4T1 and E0771 tumour bearing mice. There was no significant difference in either between the two tumour types. (Fig. 4C). Tumour volumes for each cell line are detailed in Table 2. There was no significant difference in tumour volumes for the two groups of mice and no significant relationship between tumour volume and MPI signal.

**Table 2.** Tumour volumes for mouse mammary cell lines.

| Tumour Volume (cm<sup>3</sup>) |  |  |  |  |  |  |  |  |
| --- | --- | --- | --- | --- | --- | --- | --- | --- |
| 4T1 | 0.07 | 0.01 | 0.01 | 0.03 | 0.09 | 0.18 | 0.11 | 0.15 |
| E0771 | 0.13 | 0.11 | 0.21 | 0.47 | 0.09 | 0.15 | 0.30 | 0.21 |

### Ex vivo tumour imaging

Ex vivo imaging of tumours using MPI shows presence of signal in all of the tumours in mice that were injected with iron, confirming that signal observed in vivo is from tumours (Fig. 5). There was no signal detected by MPI in the tumours from control mice.

**Figure 5.**
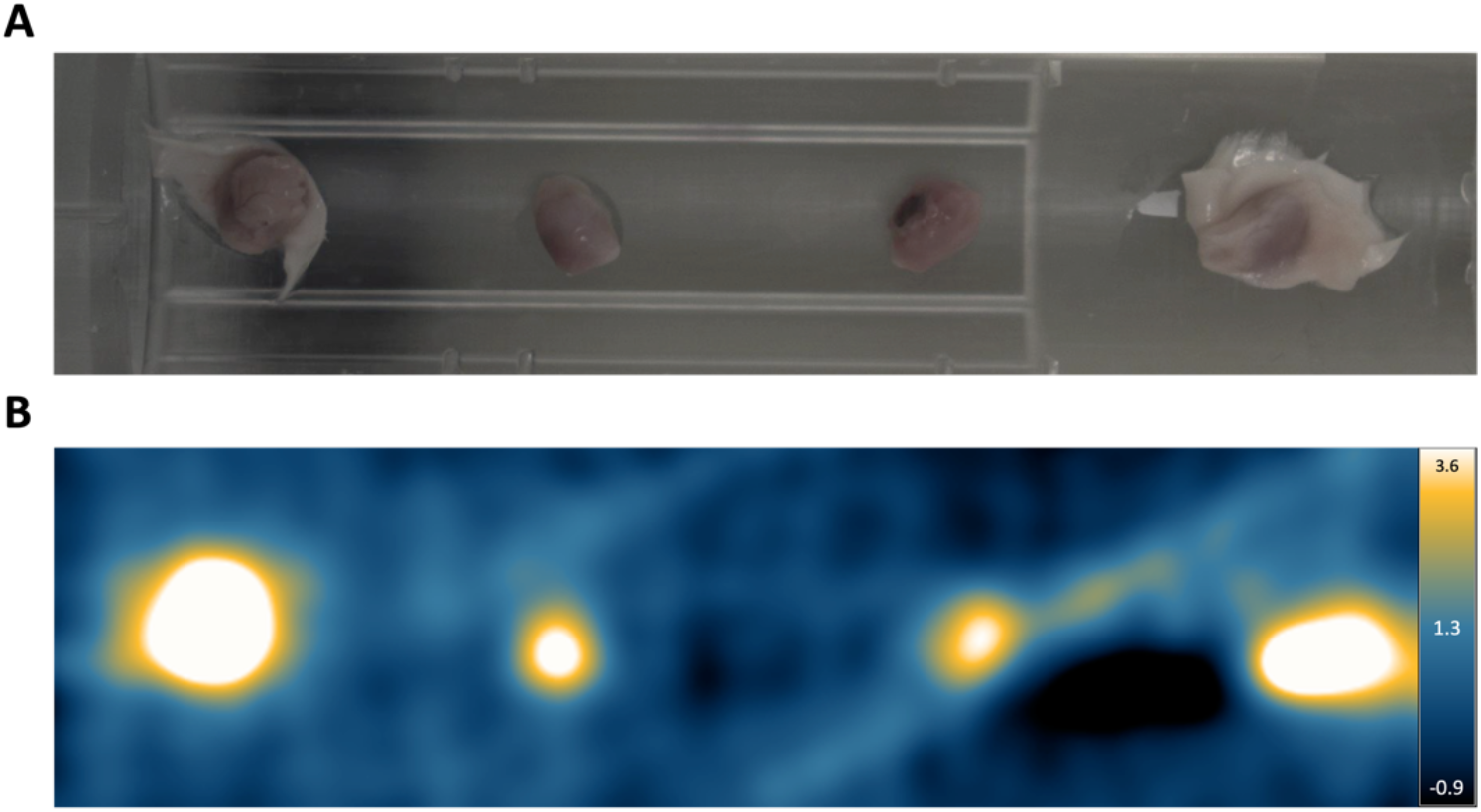
Ex vivo imaging of 4T1 tumours with MPI to determine iron presence. (A) Four representative 4T1 tumours from Balb/c mice can be seen. Tumours were placed on an empty MPI bed for imaging (2D, High sensitivity isotropic). (B) Corresponding MPI signal for each tumour can be seen, with signal detected for all tumours imaged. These results confirm the presence of SPIO in the tumours.

### Microscopy

Immunohistochemistry of tumour sections was performed to stain for CD68, a pan macrophage marker that stains for macrophages (brown, Figure 6). The staining of 4T1 tumours revealed TAMs scattered throughout the tumour. In contrast, TAMs in E0771 tumours were primarily seen on the periphery of tumour sections.

**Figure 6.**
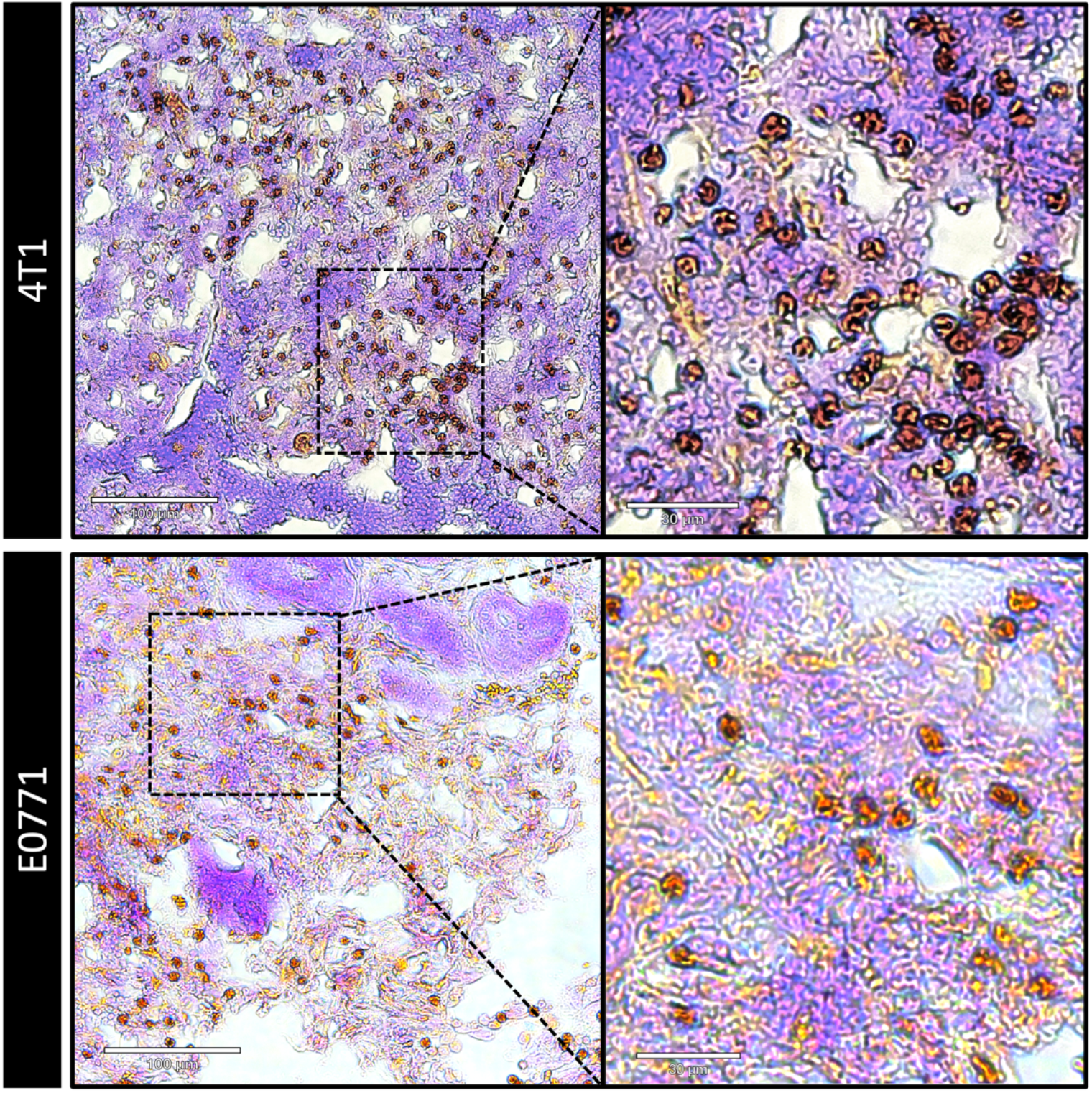
Immunohistochemistry staining of tumours showing representative CD68 positive macrophages (stained brown) for 4T1 and E0771 tumour sections.

## Discussion

Our group, and others, have been challenged by resolving proximal high and low MPI signals^19,21^. This is especially limiting for pre-clinical cell tracking experiments in animal models where a bolus of intravenously injected SPIO accumulates in the liver or when a high number of SPIO labeled cells is initially injected, and a small percentage of the cells are expected to migrate to different areas of the body. When these two signals are close to each other, their signal distributions may coalesce and prevent signal separation. This has also been referred to as “shine-through”. This is detrimental to signal quantification since ideally, only signals from the area of interest should only be included for quantification. To optimize cell tracking with MPI, this study describes the use of an advanced image reconstruction algorithm that allows for the choice of a small focused FOV to detect and quantify SPIO signal despite its proximity to a higher source of SPIO signal.

Comparing the results from the dilution series of a single sample (Figure 1) with the dilution series of two samples (Figure 2) shows the reduced dynamic range of the MPI system. In the experiments for determining the dynamic range using two samples (high iron mass + low iron mass), to visualize and quantify the lower iron concentrations, the window level had to be adjusted to the minimum and maximum signal of the lower signal; this also put the higher iron concentration within this window level, oversaturating the signal. The lower iron concentrations were quantified until S7 (780 ng Fe), at which point the lower signal could not be isolated from S1 (50.0 µg Fe). In these experiments, within a full 12 cm FOV, the native reconstruction algorithm could resolve a dynamic range of about 2^5^ = 32 for a distance of 2 cm. In Figure 5, we explored the use of a 2 cm focused small FOV centered on the low iron mass sample as a solution to the challenge of isolating weaker signal from the dominating higher signal. With the small FOV, a new reconstruction algorithm^11^ was used that produced an artifact-free image of the low iron sample without impeding signal from the higher sample. This dynamic range issue has been explored by others^22-24^. Boberg et al. proposed a two-step reconstruction algorithm which increased the dynamic range and reduced image artifacts by separating the total signal into two constituents, high and low, and using different thresholds for their respective reconstruction. Graeser et al. also proposed a type of reconstruction improvement which combined suppressing the excitation signal at its base frequency via a band-stop filter and recovering the particle signal by compensating the excitation signal in the receive chain^25^. Optimized image reconstruction is being demonstrated as a solution to low signal isolation; however, on commercially available systems this may be challenging when implementation and accessibility would come from the manufacturer.

Previously, the use of a focused small field of view (FOV) to isolate low signal regions was challenging, if not impossible, with the prescribed native reconstruction algorithm equipped on the Momentum™ MPI scanner. The regular native image reconstruction algorithm assumed that there is no signal along the edges of the FOV; if there was signal present, the values were set to zero for each line along the transmit axis^12^. This assumption caused an inverted negative image artifact when there was signal present at the FOV edge which prevented signal detection and quantification if the assumptions were not met.

We applied this approach to imaging of iron accumulation in macrophages within breast tumours in two cancer models where SPIO was administered IV, and the liver signal was the problem, and were able to isolate tumour signal in all mice and show that it can be attributed to the accumulation of iron in the tumours. Notably, the use of a focused small FOV resulted in a 50% reduction in image acquisition time.

The in vivo observation of MPI signal within tumours was validated through ex vivo imaging and IHC. We observed MPI signal in all ex vivo tumors (figure 5) and CD68 positive macrophages were detected in all tumours (figure 6). These validation steps allow us to conclude that the in vivo MPI signal is caused by SPIO within tumours and that tumours are macrophage laden. Further microscopy is required to prove that the CD68 positive macrophages contain SPIO.

We did not see a significant difference in the average MPI signal of tumours from 4T1 versus E0771 mice. There was no strong correlation between tumour size and iron content despite what has been seen in previous literature^26^. There was also no difference in average iron /mm^3^ between the two groups (data not shown). Based on previous literature we expected that the number of TAMs would be significantly higher in the more aggressive cell line, although TAMs have not been directly compared for tumours induced by the 4T1 and E0771 cell lines. This finding may be indicative of the heterogenous nature of tumour microenvironments within and between the cancer groups studied. Further work is necessary to understand this better.

One of the challenges for quantifying TAMs with MPI is estimating cell number from measurements of iron. For cell tracking studies where cells are pre-labeled with SPIO prior to their injection or transplantation a sample of labeled cells can be used to measure iron/cell (pg), the iron mass measured from the images can then be divided by this number to calculate cell number. However, when SPIO is injected IV the measurement of iron/cell is not possible. Makela et al. labeled a sample of macrophages in vitro to obtain an estimate of the in situ labeling of macrophages through an injection of IV SPIO and used this to approximate cell number^9^.

Another limitation of MPI is the lack of an anatomical image which would allow for precise localization of signals. Although we can determine that the MPI signals we observed came from the area where the tumour was implanted, we do not know if other tissues nearby have accumulated SPIO. Similarly, the relatively low resolution of MPI can limit the ability to accurately pinpoint the signal location. More recent versions of the Momentum system are equipped with a Computed Tomography (CT) scanner which mitigates this issue. MPI resolution is expected to improve as MPI tailored SPIO are created.

We chose to use SD PEG for in vivo imaging because the PEG coating is known to enhance blood circulation half-life of SPIOs^27^. Despite the PEG coating being known to reduce opsonization of SPIOs by Kupffer cells, its blood circulation half-life is comparable to Ferucarbotran, another commercially available SPIO^28^. SD PEG is not TAM-specific, and we did observe uptake in liver macrophages. There is the potential to lower liver macrophage uptake by using custom tailored particles with better blood circulation half lives^28^. Additionally, by targeting M2 macrophages specifically, as was done by Wang et al., there is the potential to preferentially target the TAMs^29^.

The primary focus of this study was to demonstrate the feasibility of using a small focused FOV in MPI to resolve and quantifying discrete sources of low signal while minimizing interference from large, nearby iron concentrations that are not of interest. We showed, for the first time, that this can be used to isolate and quantify MPI signal in tumours. This is also relevant for other in vivo MPI cell tracking research. For example, longitudinal tracking of SPIO labeled cells unwanted signal from the gastrointestinal tract can interfere with quantification of cells of interest^30^. The same approach was recently demonstrated for quantifying the migration of dendritic cells to the popliteal lymph node (small signal) after their injection into the mouse footpad (large signal)^10^. In conclusion, this work paves the way for improved in vivo preclinical MPI applications.

## Acknowledgements

Special thanks to Dr. Olivia Sehl for technical assistance and Dr. Stefaan Soenen [Katholieke Universiteit Leuven] for gifting the E0771 and 4T1 cells used in this study. We would also like to acknowledge funding from the Canadian Foundation for Innovation and the IMPAKT Facility at Western University.

## Authors Statement

Conflict of interest: Authors state no conflict of interest. Ethical approval: The research related to animal use complies with all the relevant national regulations, institutional policies and was performed in accordance with the standards of the Canadian Council on Animal Care, under an approved protocol by the Animal Use Subcommittee of Western University’s Council on Animal Care.

